## Supplementary file for "A hepatocyte-specific transcriptional program driven by Rela and Stat3 exacerbates experimental colitis in mice by modulating bile synthesis"

### Supplementary Figures and Tables

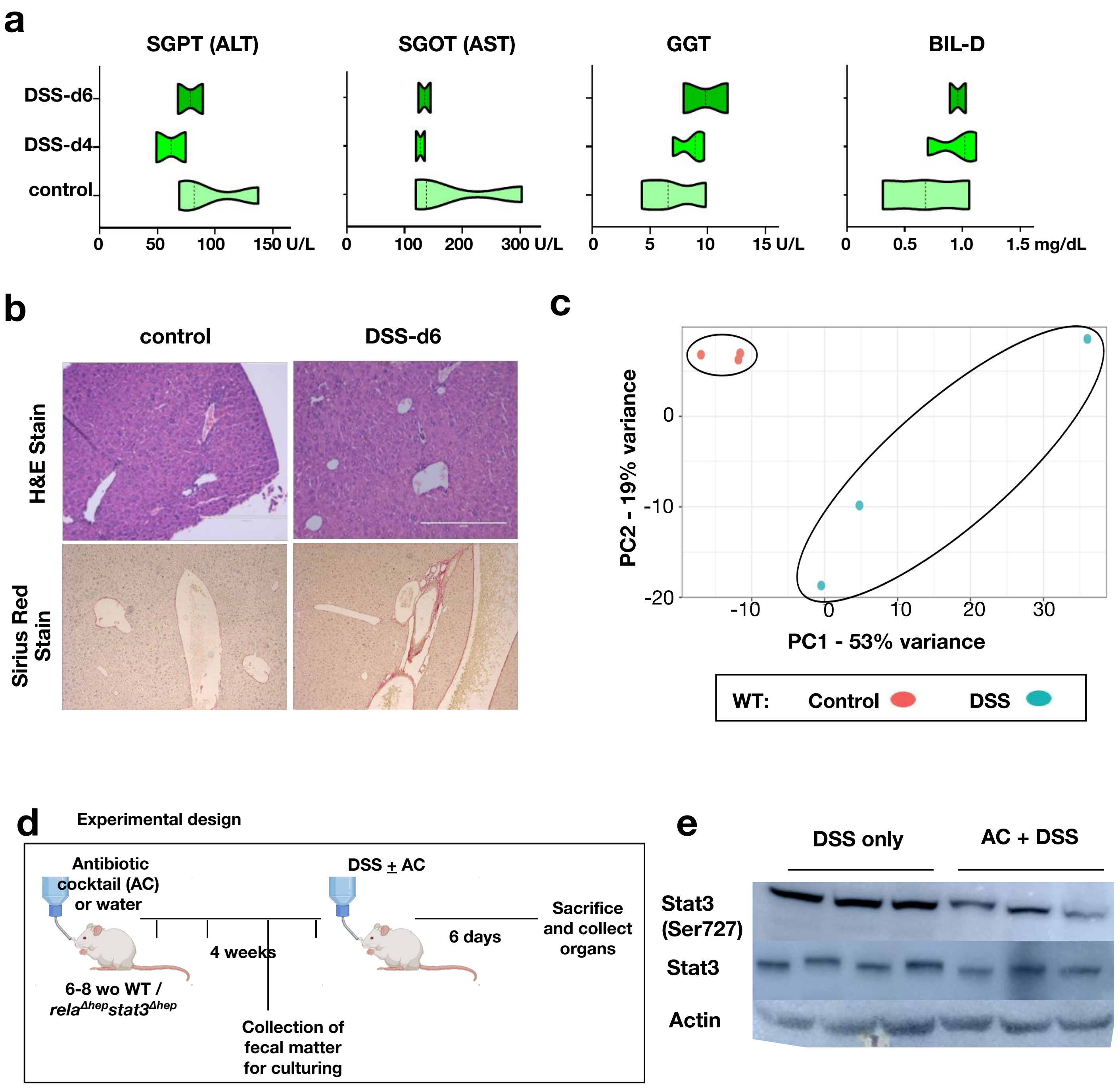


#### Supplementary Figure S1

**Figure S1: (a)** Violin plot comparing the clinical parameters in serum of untreated and DSS-treated wild type mice (n=3) **(b)** Liver sections from untreated and DSS-treated wild type mice were examined for histological features involving H&E staining [upper panel] and Sirius red staining [lower panel]. Data were obtained in 10X magnification and represent three experimental replicates; two fields per section and a total of three sections from each set were examined. **(c)** PCA plot illustrating the hepatic transcriptome, identified through global RNA-seq analyses, of untreated or DSS treated wild type mice (n=3). **(d)** Schematic description of experimental setup for gut sterilization by antibiotic treatment. **(e)** Western blot revealing the abundance of total Stat3 and it’s phosphorylated functionally active forms p-Stat3 (Ser727), in the liver extracts prepared from DSS-treated wild type C57BL/6 mice with and without a prior antibiotic treatment for four weeks.

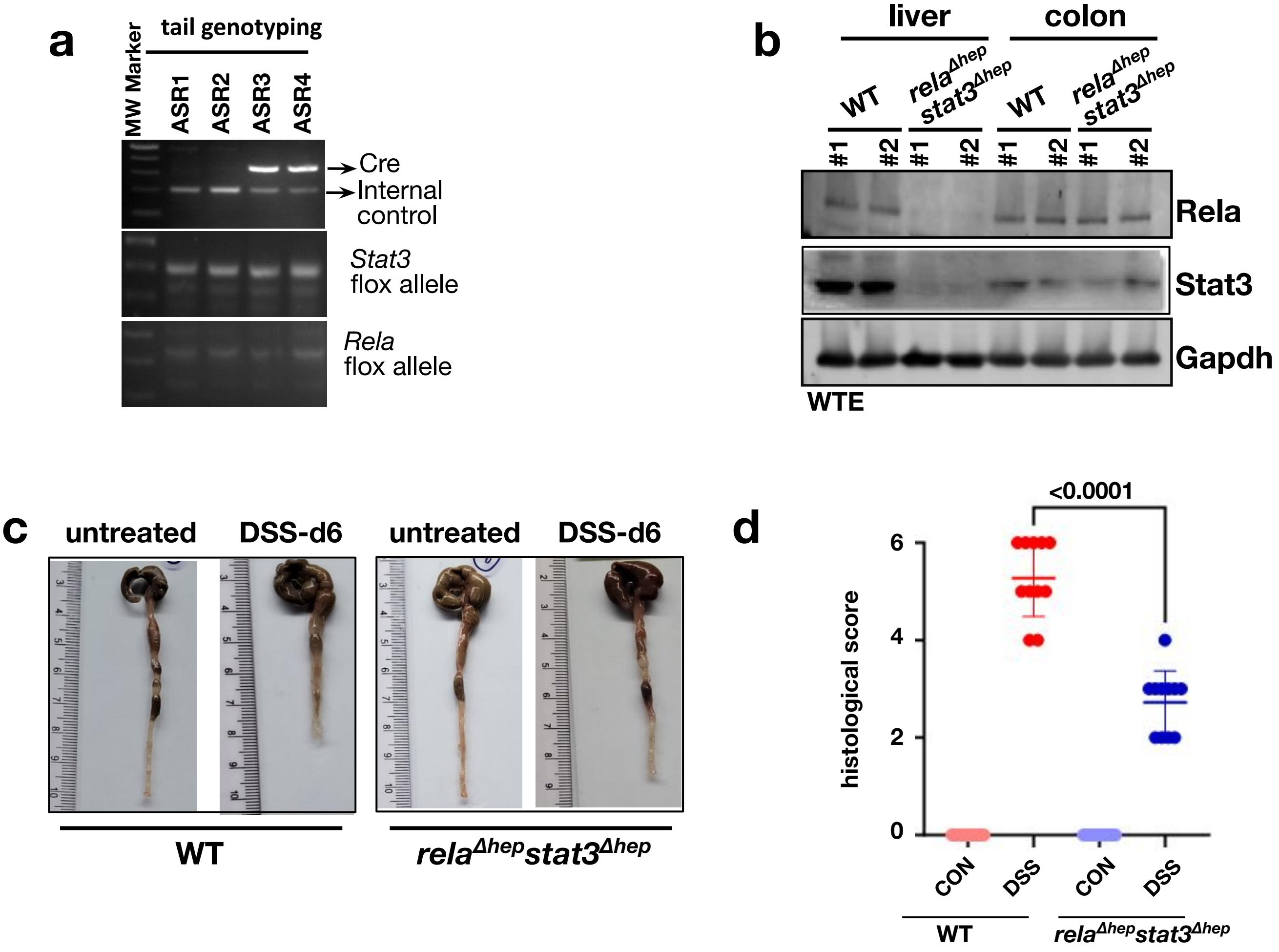


**Figure S2:** Characterization of mice strain by **(a)** genotyping and **(b)** western blotting; WTE - whole tissue extract. **(c)** Representative images of colon from untreated and DSS-treated wild type and *rela*$^{\Delta hep}$*stat3*$^{\Delta hep}$ mice. **(d)** Dot plot representing the histological score for H&E stained colon sections in figure 2C, each dot represent the score for each field analyzed.

### Figure S2

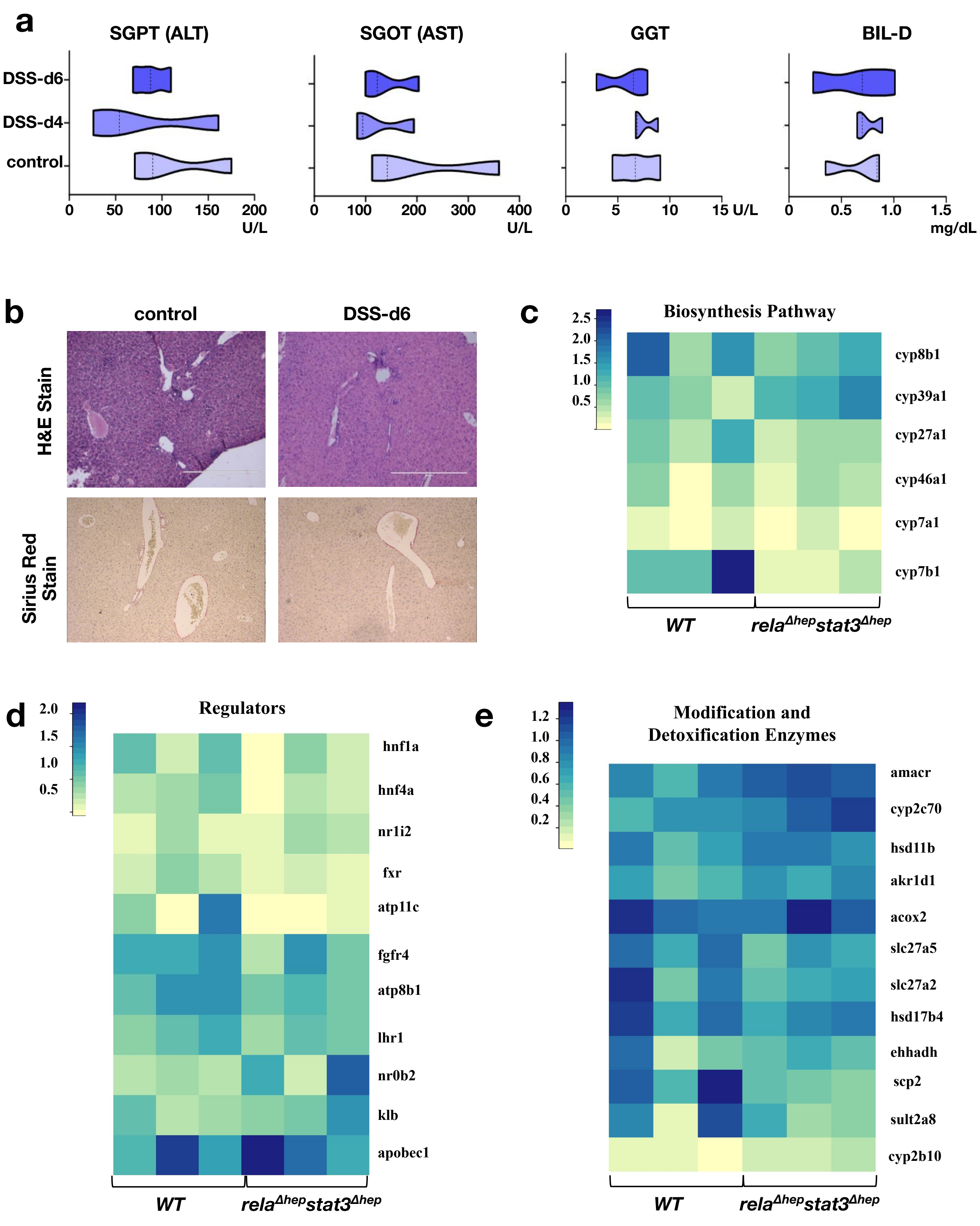
a
SGPT (ALT)
SGOT (AST)
GGT
BIL-D
DSS-d6
DSS-d4
control
0
50
100
150
200
U/L
0
100
200
300
400
U/L
0
5
10
15 U/L
0
0.5
1.0
1.5
mg/dL
b
control
DSS-d6
H&E Stain
Sirius Red Stain
c
Biosynthesis Pathway
2.5
2.0
1.5
1.0
0.5
cyp8b1
cyp39a1
cyp27a1
cyp46a1
cyp7a1
cyp7b1
WT
relaΔhepstat3Δhep
d
Regulators
2.0
1.5
1.0
0.5
hnf1a
hnf4a
nr1i2
fxr
atp11c
fgfr4
atp8b1
lhr1
nr0b2
klb
apobec1
WT
relaΔhepstat3Δhep
e
Modification and
Detoxification Enzymes
1.2
1.0
0.8
0.6
0.4
0.2
amacr
cyp2c70
hsd11b
akr1d1
acox2
slc27a5
slc27a2
hsd17b4
ehhadh
scp2
sult2a8
cyp2b10
WT
relaΔhepstat3Δhep

### Figure S3

**Figure S3:** **(a)** Violin plot comparing the clinical parameters in serum of untreated and DSS-treated *rela*$^{\Delta hep}$*stat3*$^{\Delta hep}$ mice (n=3) **(b)** Liver sections from untreated and DSS-treated *rela*$^{\Delta hep}$*stat3*$^{\Delta hep}$ mice were examined for histological features involving H&E staining [upper panel] and Sirius red staining [lower panel]. Data were obtained in 10X magnification and represent three experimental replicates; two fields per section and a total of three sections from each set were examined. Heatmap represents relative transcript abundance of **(c)** biosynthesis pathway **(d)** regulators of bile acid metabolic pathways and **(e)** modification enzymes of bile acid metabolic pathway of indicated genotypes obtained from the RNA-seq experiment of three biological replicates.

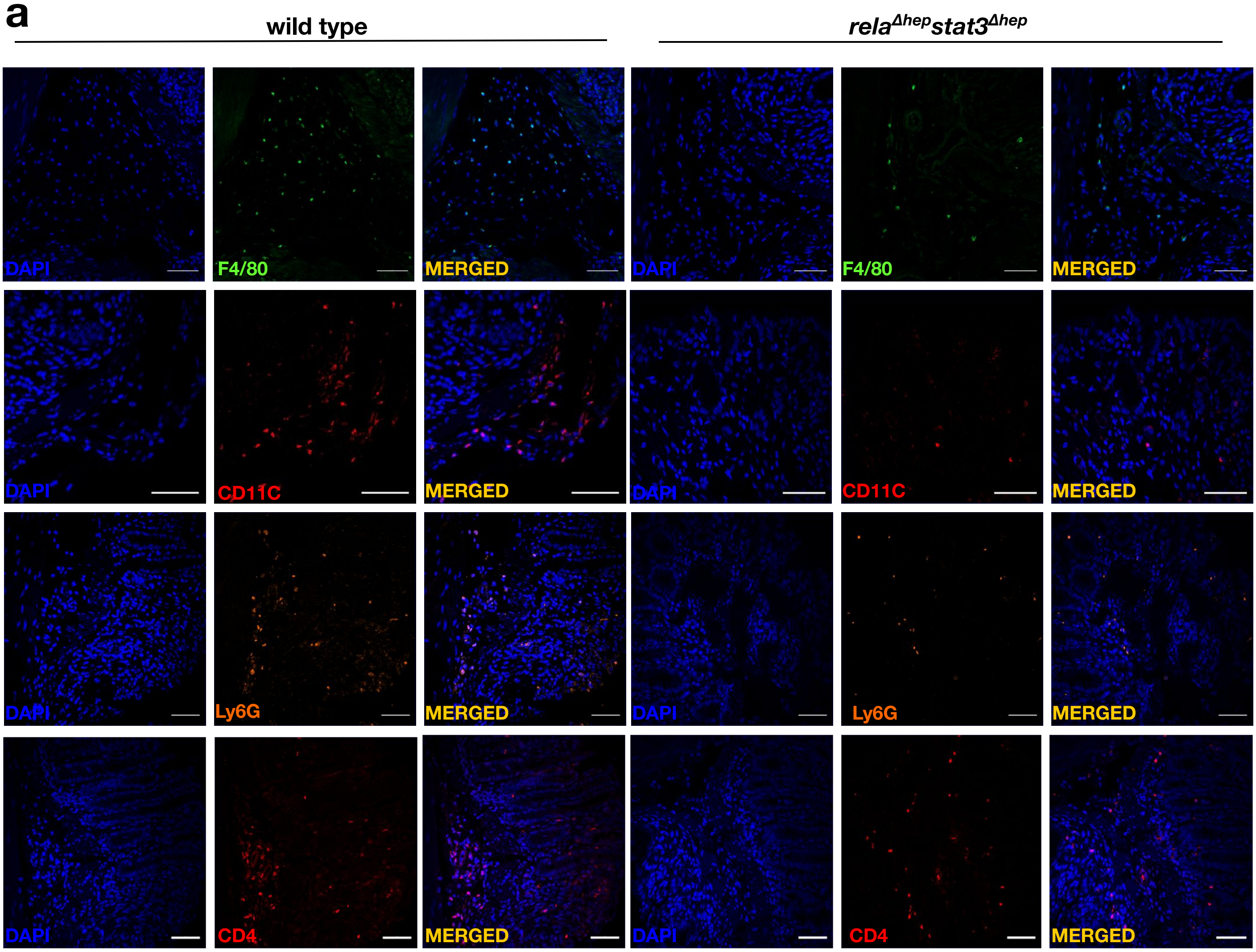


**Figure S4: (a)** Representative images of colon tissue stained with indicated immune cell markers, DAPI and merged section in the DSS-treated wild type and *rela*$^{\Delta hep}$*stat3*$^{\Delta hep}$ mice. Scale represents is 50 µm.

### Figure S4

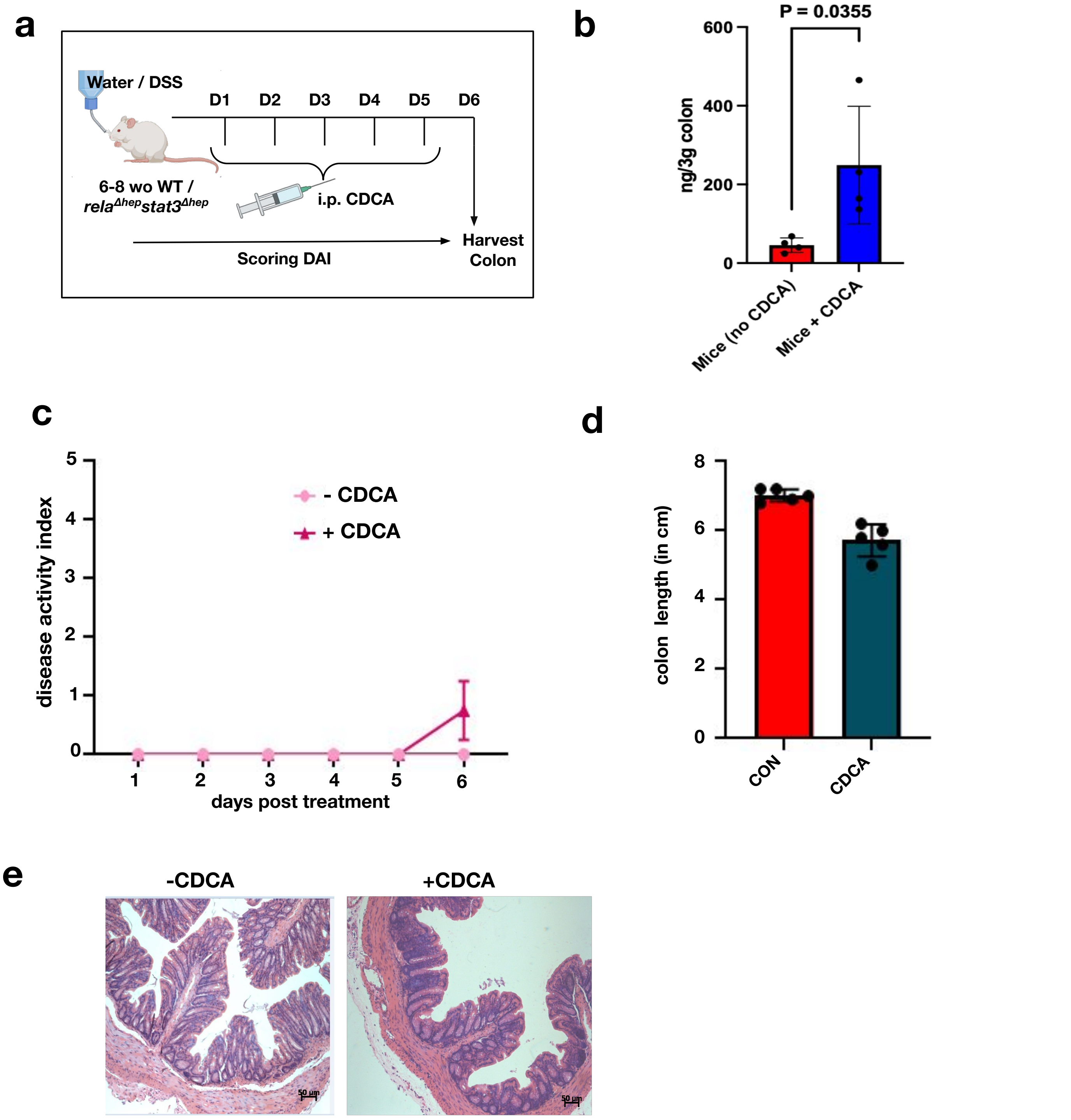


**Figure S5: (a)** Schematic description of the experimental design describing the course of DSS treatment and CDCA supplementation. **(b)** Quantification of CDCA in the colon of mice, that received intraperitoneal injection of either CDCA or DMSO (control), through targeted LC-MS experiments. **(c)** Line plot charting the disease activity in a time course of *rela*$^{\Delta hep}$*stat3*$^{\Delta hep}$ mice subjected to daily supplementation of 10 mg/kg CDCA. Mice devoid of CDCA supplementation were treated with DMSO as controls. **(d)** Bar plot comparing the colon length of *rela*$^{\Delta hep}$*stat3*$^{\Delta hep}$ mice subjected to CDCA supplementation. **(e)** Colon sections from *rela*$^{\Delta hep}$*stat3*$^{\Delta hep}$ mice supplemented with CDCA were examined by H&E staining.

### Figure S5

**Supplementary methods:**

**Histological Scoring:** Three experimental replicates was used for histological scoring, two fields per section and a total of three sections from each set were examined. Histological injury was assessed by a combined score of inflammatory cell infiltration (score 0–3) and mucosal damage (score 0–3) as previously described in *Ren Y et al.* 2019 (10.1038/s41598-019-53305-z).

**Antibiotic treatment:**

Gut sterilization was achieved by administration of an antibiotic cocktail: ampicillin 1g/l, neomycin 1g/l, metronidazole 0.25g/l, and vancomycin 0.5g/l as described in *Hernández-Chirlaque C et al.* 2016. Antibiotic treatment was applied for 4 week before the start of DSS-treatment and was maintained until the end of the experiment. Depletion of gut microbiota was confirmed by conventional bacterial enumeration studies from the faeces of mice before and during the antibiotic treatment.

**Table S1: List of antibodies**

| Antibody | Source | Identifier | RRID |
| --- | --- | --- | --- |
| PE rat anti-mouse Ly-6G | BD Biosciences | 551461 | AB_394208 |
| FITC anti-mouse F4/80 | BioLegend | 157309 | AB_2876535 |
| PE anti-mouse CD11c | BioLegend | 117308 | AB_313777 |
| PE anti-mouse CD4 | BioLegend | 100408 | AB_312693 |
| STAT3 mouse mAb | Thermo Fisher Scientific | MA1-13042 | AB_10985240 |
| RelA rabbit pAb | Santa Cruz Biotechnology | sc372 | AB_632037 |
| Phospho-Stat3 (Ser727) Rabbit mAb | Cell Signaling Technology | 34911 | AB_2737598 |
| Phospho-Stat3 (Ser727) Mouse mAb | Cell Signaling Technology | 9136 | AB_331755 |
| Phospho-Stat3 (Tyr705) Rabbit mAb | Cell Signaling Technology | 9145 | AB_2491009 |
| Phospho-NF-κB p65 (Ser536) pAb | Cell Signaling Technology | 3031 | AB_330559 |
| GAPDH rabbit mAb | Cell Signaling Technology | 2118 | AB_561053 |
| β-Actin rabbit pAb | Cell Signaling Technology | 4967 | AB_330288 |
| Goat anti-Rabbit IgG (H+L) Secondary Antibody Alexa Fluor Plus 555 | Thermo Fisher Scientific | A32732 | AB_2633281 |

**Table S2: List of mice strain**

| Mice strain | Source | Identifier |
|---|---|---|
| C57BL/6 WT mice | available at the Small Animal Facility in NII | N/A |
| *Alb_Cre rela$^{-/-}$ stat3$^{-/-}$* mice | gift from Dr. Lee Quinton (Boston University) | N/A |
| *rela$^{fl/fl}$* stat3$^{fl/fl}$ mice | gift from Dr. Lee Quinton (Boston University) | N/A |
| *Alb_Cre rela$^{-/-}$* mice | gift from Dr. Lee Quinton (Boston University) | N/A |
| *rela$^{fl/fl}$* mice | gift from Dr. Lee Quinton (Boston University) | N/A |
| *Alb_Cre stat3$^{-/-}$* mice | gift from Dr. Lee Quinton (Boston University) | N/A |
| stat3$^{fl/fl}$ mice | gift from Dr. Lee Quinton (Boston University) | N/A |

**Table S3: List of genotyping primers**

| Target gene | Sense strand | Antisense strand |
|---|---|---|
| Cre | 5'-GGTGAACGTGCAAAACAGGCTC-3' | 5'-AAAACAGGTAGTTATTCGGATCATCAGC-3' |
| Tcrd (Internal control) | 5'-CAAATGTTGCTTGTCTGGTG-3' | 5'-GTCAGTCGAGTGCACAGTTT-3' |
| Floxed Stat3 | 5'-CCTGAAGACCAAGTTCATCTGTGTTGAC-3' | 5'-CACACAAGCCATCAAACTCTGGTCTCC-3' |
| Floxed Rela | 5'-GAGCGCATGCCTAGCACCAG-3' | 5'-GTGCACTGCATGCGTGCAG-3' |

**Table S4: List of primers**

| Target gene | Sense strand | Antisense strand |
|---|---|---|
| Tight junction protein 1 (tjp1) | 5’-GCTTTAGCGAACAGAAGGAGC-3’ | 5’-TTCATTTTTCCGAGACTTCACCA-3’ |
| Occludin (ocln) | 5’-TGAAAGTCCACCTCCTTACAGA-3’ | 5’-CCGGATAAAAAGAGTACGCTGG-3’ |
| Mucin 2 (muc2) | 5’-AGGGCTCGGAACTCCAGAAA-3’ | 5’-CCAGGGAATCGGTAGACATCG-3’ |
| Trefoil factor 3 (tff3) | 5’-TTGCTGGGTCCTCTGGGATAG-3’ | 5’-TACACTGCTCCGATGTGACAG-3’ |
| Interleukin-1β (il1β) | 5’-CATCCCATGAGTCACAGAGGATG-3’ | 5’-ACCTTCCAGGATGAGGACATGAG-3’ |
| Tumor necrosis factor α (tnfα) | 5’-CTGAACTTCGGGGTGATCGG-3’ | 5’-GGCTTGTCACTCGAATTTTGAGA-5’ |
| Interleukin-6 (il6) | 5’-CCCCAATTTCCAATGCTCTCC-3’ | 5’-GGATGGTGTTGGTCCTTAGCC-3’ |
| gapdh | 5’-AGGTCGGTGTGAACGGATT-3’ | 5’-AATCTCCACTTTGCCACTGC-3’ |
| cyp7a1 | 5’-GCTGTGGTAGTGAGCTGTTG-3’ | 5’-GTTGTCCAAAGGAGGTTCACC-3’ |
| cyp8b1 | 5’-CCTCTGGACAAGGGTTTTGTG-3’ | 5’-GCACCGTGAAGACATCCCC-3’ |
| cyp27a1 | 5’-AGGGCAAGTACCCAATAAGAGA-3’ | 5’-TCGTTTAAGGCATCCGTGTAGA-3’ |
| cyp7b1 | 5’-TCCTGGCTGAACTCTTCTGC-3’ | 5’-CCAGACCATATTGGCCCGTA-3’ |

**Table S5: List of reagents**

| Reagent | Source | Identifier |
|---|---|---|
| **Chemicals and Peptides** | | |
| DSS (40,000 MW) | Sigma-Aldrich | 42867 |
| FITC-Dextran (MW 3000 - 5000) | Sigma-Aldrich | 60842-46-8 |
| Immobilon Forte Western HRP Substrate | Millipore | WBLUF0500 |
| Amersham™ Hybond® P Western blotting membranes, PVDF | Merck | 10600023 |
| 1.0 mm Zirconia/Silica | BioSpec Product | 11079110z |
| Chenodeoxycholic acid | Sigma-Aldrich | 474-25-9 |
| DAPI | Sigma-Aldrich | D9542 |
| PowerUp SYBR | Thermo Fisher Scientific | A25742 |
| Fluorosheild | Sigma-Aldrich | F6182 |
| **Commercial Kits** | | |
| NucleoSpin RNA | Macherey-Nagel | 74106 |
| Primescript 1$^{st}$ strand cDNA synthesis kit | Takara Bio | 6110A |

**Table S6: Control patient demography**

| Sample code | Sample type | Age (years) | Gender (1=Male) | BMI |
|---|---|---|---|---|
| 01-C-F | Control | 50 | 2 | 24.6 |
| 02-C-F | Control | 35 | 1 | 21.8 |
| 03-C-F | Control | 50 | 1 | 16.9 |
| 04-C-F | Control | 27 | 2 | 17.7 |
| 05-C-F | Control | 42 | 2 | 23.4 |
| 06-C-F | Control | 36 | 1 | 25 |
| 07-C-F | Control | 54 | 1 | 21.3 |
| 08-C-F | Control | 52 | 2 | 23.3 |
| 10-C-F | Control | 19 | 1 | 21.17 |
| 12-C-F | Control | 30 | 1 | 17.8 |
| 13-C-F | Control | 28 | 1 | 21.3 |
| 14-C-F | Control | 59 | 2 | 20.2 |
| 16-C-F | Control | 32 | 1 | 23.5 |

**Table S7: UC patient demography**

| Sample code | Sample type | Age (years) | Gender (1=Male) | BMI |
|---|---|---|---|---|
| 01-A-F | UC | 50 | 2 | 25 |
| 02-A-F | UC | 44 | 1 | 24 |
| 03-A-F | UC | 29 | 1 | 16 |
| 04-A-F | UC | 25 | 1 | 15 |
| 05-A-F | UC | 32 | 1 | 18 |
| 06-A-F | UC | 50 | 2 | 23 |
| 07-A-F | UC | 24 | 1 | 17 |
| 08-A-F | UC | 50 | 1 | 26 |
| 09-A-F | UC | 27 | 2 | 22 |
| 10-A-F | UC | 35 | 1 | 20 |
| 11-A-F | UC | 22 | 2 | 18 |
| 12-A-F | UC | 25 | 2 | 23 |
| 13-A-F | UC | 30 | 1 | 29 |
| 14-A-F | UC | 26 | 1 | 17 |
| 15-A-F | UC | 25 | 2 | 17 |
| 17-A-F | UC | 24 | 2 | 29 |
| 18-A-F | UC | 26 | 1 | 21 |
| 19-A-F | UC | 45 | 2 | 21 |
| 20-A-F | UC | 23 | 1 | 18 |
| 21-A-F | UC | 24 | 2 | 17 |
| 22-A-F | UC | 26 | 1 | 17 |
| 23-A-F | UC | 50 | 2 | 37 |
| 24-A-F | UC | 43 | 1 | 25 |
| 25-A-F | UC | 30 | 2 | 23 |
| 26-A-F | UC | 31 | 1 | 16 |
| 27-A-F | UC | 32 | 1 | 19 |
| 28-A-F | UC | 26 | 1 | 23 |
